# A targeted mutational strategy aiding generating antisense RNA to knockdown the *Ehrlichia chaffeensis* p28-outer membrane protein 19 expression

**DOI:** 10.64898/2026.01.05.697664

**Authors:** Xishuai Tong, Dominica Ferm, Huitao Liu, Ying Wang, Chandramouli Kondethimmanahalli, Roman R. Ganta

## Abstract

Obligate intracellular pathogenic bacteria belonging to the order Rickettsiales include several important emerging pathogens causing major health and economic impact to people, companion animals, and agricultural animals. Despite some recent progress, the lack of well-established genetic manipulation methods for diverse research applications remains a challenge. We recently reported the establishment of targeted mutagenesis methods to disrupt genes in *Ehrlichia* and *Anaplasma* species. Many essential genes in *Ehrlichia chaffeensis* are likely refractory to targeted mutagenesis, thus we developed a novel targeted mutational approach leading to the expression of antisense RNA to facilitate the knockdown of p28-Omp19 protein expression from *ECH_1143.* This gene was selected as its encoded protein is among the highly immunogenic proteins of *E. chaffeensis* and is likely essential for the pathogen. This method involved introducing a mutation at a distal genomic location within the *E. chaffeensis* genome to allow for generation of a 209 nucleotide-long antisense RNA segment complementary to *ECH_1143* coding mRNA from the same gene promoter which was duplicated as part of the mutagenesis. The mutational strategy was designed to retain the surrounding genomic regions unaltered. The antisense knockdown version of *E. chaffeensis* resulted in a reduction of p28-Omp19 expression when compared to wild-type *E. chaffeensis* during its replication in a macrophage cell line, where the gene expression is known to occur. We anticipate that the antisense mutational strategy will be broadly applicable to facilitate investigating essential genes of obligate intracellular bacterial pathogens.

## INTRODUCTION

Obligate intracellular pathogenic bacteria are responsible for causing diseases in millions of humans and several vertebrates worldwide (1, 2). A well-developed targeted mutagenesis system supporting research on obligate intracellular bacteria belonging to the order Rickettsiales within the genera *Ehrlichia, Anaplasma, Neorickettsia, Rickettsia,* and *Orientia* continue to be a major limiting factor for research progress. Several pathogens that belong to these genera are responsible for causing emerging and reemerging tick-borne diseases in humans and various vertebrate animals (1, 3, 4). The pathogens have extensive genome reductions (5–7) and retained predominantly genes critical primarily for intracellular growth adaptation which creates a unique challenge for defining molecular genetics broadly applicable for diverse research applications. Furthermore, *Anaplasmataceae* family bacteria naturally lack plasmids, thus limiting the ability to use a plasmid in manipulating the bacterial gene expression for pathogens of the genera *Anaplasma, Ehrlichia* and *Neorickettsia*. Despite recent progress in using Himar1 random mutagenesis and the development of targeted mutations in Rickettsiales, the probability of generating targeted mutations that cause complete loss of a gene function remains very low (8–12). To date, only a limited number of targeted mutations are described in defining the functional characterization in rickettsial pathogens including the genera *Anaplasma*, *Ehrlichia* and *Rickettsia* (8–10, 13–15). Thus, it is evident that additional methodological advances are necessary in developing molecular tools for promoting research on rickettsial pathogens.

Recently, we reported the development of targeted mutagenesis methods that work well for two *Anaplasmataceae* family pathogens; *Anaplasma marginale* and *Ehrlichia chaffeensis* (8–11). We also demonstrated functional disruption and restoration mutations in *E. chaffeensis* (11). We reasoned that due to the pathogens reduced genome(s), the majority of the genes are conceivably essential for the bacterial intracellular adaptation, and thus intractable to mutations resulting in a complete loss of functional disruption or ablation. Therefore, we opted to develop a targeted mutagenesis strategy to interfere with the translation of messenger RNA (mRNA) by simultaneously expressing a portion of the coding sequence as an antisense RNA (asRNA) leading to the complementary binding with sense mRNA and reducing protein expression.

Typically, asRNA molecules interact with sense mRNA through complementary base pairing to block translation and downregulate a gene expression (16). This mechanism of gene regulation is documented across diverse organisms including those belonging to archaea and eukaryotes (17–20). Exogenous synthetic antisense oligonucleotide (ASO) production can serve as a powerful tool for studying transient protein knockdown at the post-transcriptional level by hybridizing with the sense mRNA, leading to either mRNA degradation through a ribonuclease-mediated cleavage or steric hindrance of translation by blocking the interaction of mRNA with ribosomes (21, 22). The use of exogenous synthetic ASO is not ideal as it is typically short-lived and is highly susceptible to nuclease degradation, resulting a possible limited transient suppression of the target mRNA translation (21). Furthermore, application of exogenous synthetic ASO transfection can be an added challenge for investigations focused on obligate intracellular bacterial organisms, such as for Rickettsiales due to their intracellular residence within a host cell phagosome or within the cytoplasmic space. A plasmid-based expression of constitutive asRNA production has the potential to provide a continuous supply of asRNA, allowing rapid downregulation of a targeted gene expression by inhibiting translation (23, 24). Although our previous work has shown that uptake of genetic material by these organisms is possible (10, 11, 25), the use of a plasmid-based knockdown is not possible for use in Anaplasmataceae family pathogens, such as *E. chaffeensis*, because they naturally lack endogenous plasmids (26).

In the current study, we developed an innovative targeted mutagenesis approach in *E. chaffeensis* to facilitate expressing specific asRNA to inhibit protein synthesis from *ECH_1143* causing the knockdown of its encoded protein. We selected this gene as its encoded protein, p28-Omp19, is one of the most abundantly expressed immunogenic proteins of *E. chaffeensis* (27). We created an insertion mutation at a remote genomic location to produce the *trans* acting asRNA that is also made from the duplicated same gene (*ECH_1143*) promoter to reduce p28-Omp19 protein expression. Furthermore, the mutational strategy was carefully designed to restore the genomic sequences surrounding the mutational site as unaltered. The data presented in this manuscript demonstrate that the asRNA generation by the unique mutational strategy is ideal for investigating genes critical for the bacterial replication *in vitro*.

## RESULTS

### Construction of a recombinant plasmid aided in generating asRNA mutation in the *E. chaffeensis* genome

We selected *E. chaffeensis ECH_1143* for generating an asRNA mutation for its protein expression knockdown because it is one of the 22 gene paralogs encoding for the abundantly expressed and highly immunogenic protein, p28-Omp19 (28–31). A recombinant plasmid with pGGA plasmid backbone was created by following standard molecular cloning methods to inserting multiple segments in the following order (Fig 1): 1) The left homology arm (L) spanning part of *ECH_0230* coding region and the entire noncoding region located downstream to *ECH_0230*; 2) *E. chaffeensis tuf* gene promoter (8) (T); 3) mCherry gene coding sequence (11) (M); 4) *E. chaffeensis* codon optimized gentamycin resistance gene coding sequence (11) (G); 5) the entire predicted promoter segment located upstream to *ECH_1143* and the first 209 base pair segment of the *ECH_1143* cloned in reverse orientation (AS); 6) the right homology arm (R) spanning the entire downstream noncoding region of *ECH_0230* and a 3’ end segment of *ECH_0232* coding region. The duplication of the entire noncoding region downstream of *ECH_0230* was necessary to restore the *E. chaffeensis* genomic sequences as unaltered during mutation development. We selected the mutation insertion site based on our previous demonstration of generating a mutation near to this location (8). Linear fragments spanning from L to R along with the rest of the inserted segments generated by PCR and 3 μg of the purified amplicon was then used to electroporate ISE6 tick cell culture-derived, *E. chaffeensis* organisms and subsequently reinfected to ISE6 tick cells (11). Cultures expressing mCherry and resisting to gentamicin were observed after about three weeks of *in vitro* growth at 34°C. Schematic view of the asRNA generation and its anticipated interaction with the sense mRNA of *ECH_1143* is depicted in Fig 2A. Two different PCRs were performed to confirm the mutational generation besides resisting to gentamycin in the culture media and testing positive for mCherry protein expression: PCR I using primers annealing upstream and downstream to the insertion site (primers P1 and P3) yielded the expected amplicon of 4.77 kb with the mutant genomic DNA as the template, whereas wild-type *E. chaffeensis* genomic DNA yielded the smaller estimated 2.07 kb amplicon (Fig 2B left panel). Similarly, a second PCR (PCR II) targeting the insertion region and genomic region upstream to insertion site generated the expected amplicon (2.18 kb) only with the mutant genomic DNA as the template, but not with wild-type genomic DNA (Fig 2B right panel). Further, whole genome sequence analysis was performed independently to confirm the presence of the inserted sequences at the desired target site (Fig 2C).

**Figure 1.**
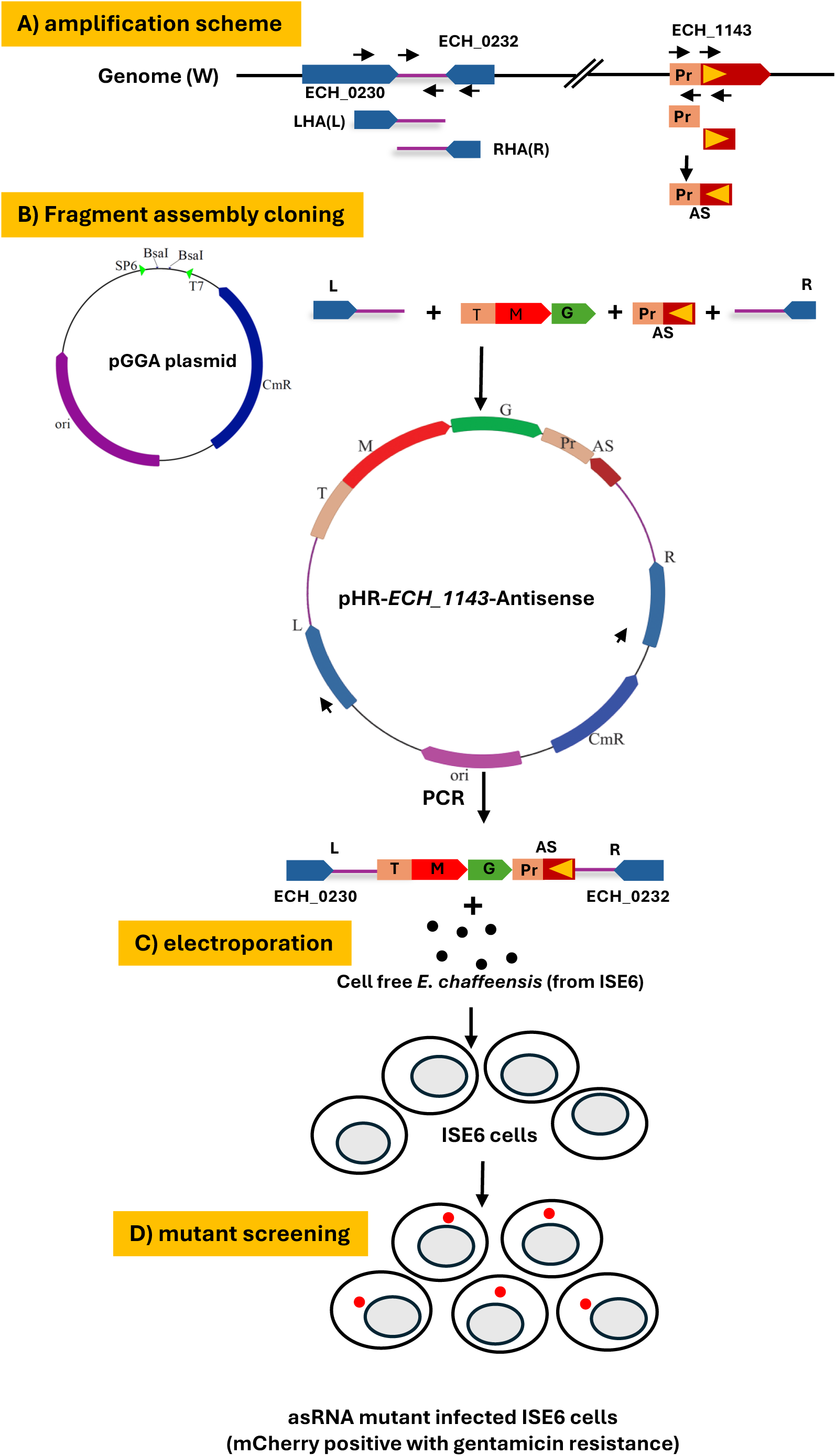
Schematic overview of the *E. chaffeensis ECH_1143* antisense construct generation and its use in creating the asRNA mutation. **A) The amplification scheme** for the left and homology arms (L and R, respectively) included in generating the same segment of 3’ noncoding region of *ECH_0230* to be positioned at the 3’ and 5’ of the amplified segments, respectively. The *ECH_1143* promoter segment and the first 209 base pairs of the ORF were amplified separately and combined with selecting the ligated ORF segment reverse orientation relative to the promoter segment. Tuf (T), mCherry ORF and gentamycin resistance cassette were obtained from a previously prepared recombinant construct (ref). B) **Fragment assembly cloning** of all the fragments into pGGA plasmid in the chosen order was accomplished using a Golden Gateway assembly cloning kit to generate the final recombinant plasmid; pHR-*ECH_1143*-Antisense. **C) Electroporation.** The mutagenesis segment spanning from L to R was generated by PCR and used to transform ISE6 tick cell culture derived *E. chaffeensis* organisms. **D) Mutant screening** to select the mutated bacteria in the tick cells was accomplished by screening for the cultures expressing mCherry and resisting to gentamicin.

**Figure 2.**
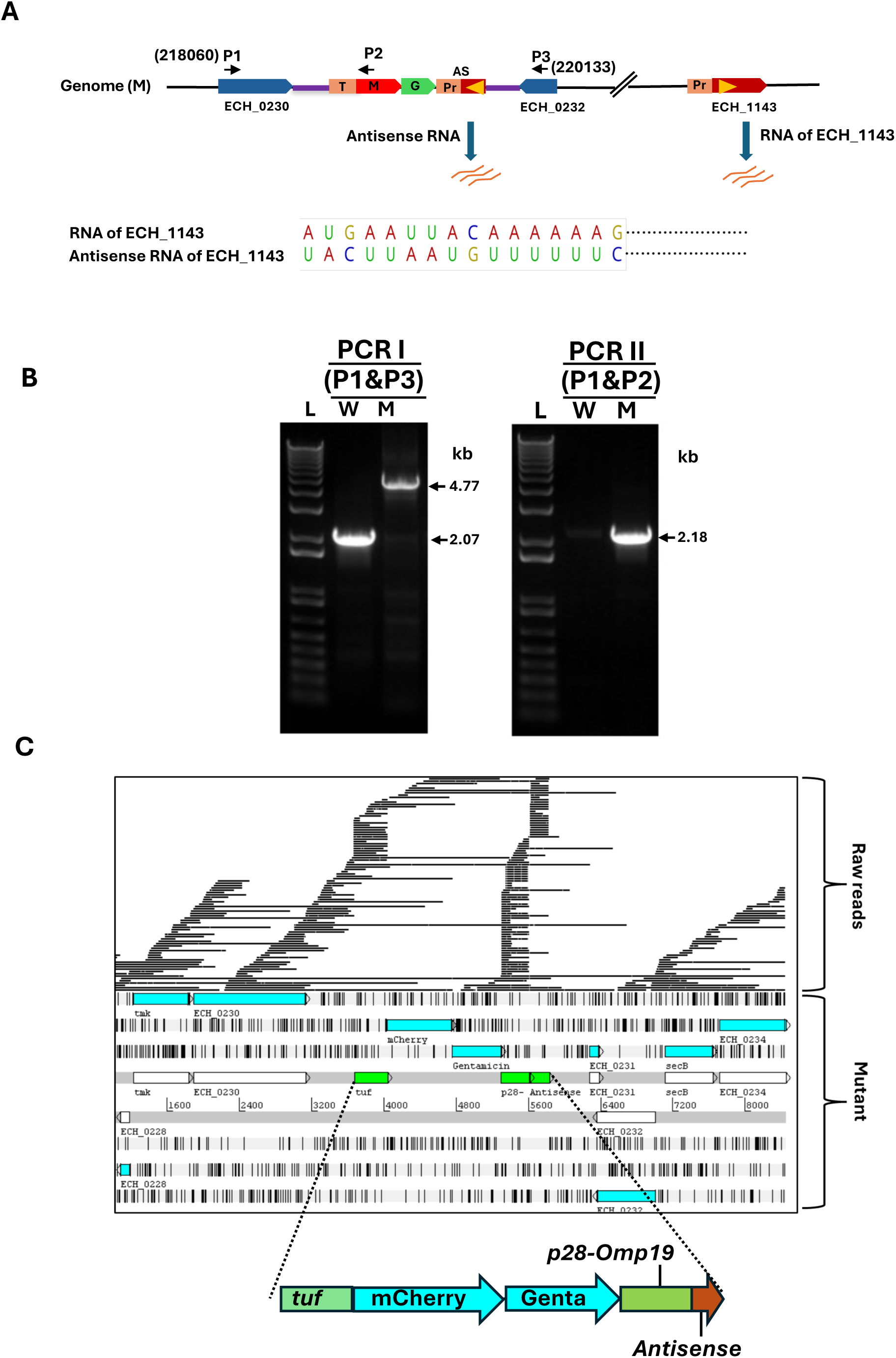
Confirmation of the mutant *E. chaffeensis* by PCR and whole genome sequencing. **A) Schematic view** of the duplex RNA strand formation of asRNA with mRNA of *ECH1143.* **B) PCR analysis** confirming the mutation generation. Primers annealing upstream (P1) and downstream (P2) of the mutation insertion region (genomic coordinates identified) were used to amplify DNA segments by PCR I which were then resolved on an agarose gel; the expected larger 4.77 kb product was evident from the mutant genomic DNA (M) compared to the smaller 2.07 kb from wild-type genomic DNA (W). (L, 1 kb plus DNA ladder). Similarly, PCR II using primers P1 and P2 targeted to insertion region and genomic region upstream to insertion region yielded the expected amplicon of 2.18 kb only from the mutant. C) Nanopore sequencing reads were used to assemble the genome of the antisense mutant. The insertion segment was identified at the anticipated region within the *E. chaffeensis* genome.

### Antisense mutation resulted in a reduction of p28-Omp19 protein expression in *E. chaffeensis* cultured in the macrophage cell line

#### Confocal microscopy assessment revealed decreased p28-Omp19 protein expression in macrophage cultures

We assessed the p28-Omp19 expression by confocal microscopy for the asRNA mutant for the macrophage and tick cell cultures harvested when infectivity of the cultures was about 80% and compared with wild-type when the organisms were cultured in macrophage and tick cells; a p28-Omp19-specific monoclonal antibody, mAb 18.1 (32) was used for this experiment (Fig 3). The mutant *E. chaffeensis* exhibited constitutive expression of mCherry, and a considerable reduction in p28-Omp19 expression compared to the wild-type in the canine macrophage line (DH82) (Fig 3A). Consistent with prior published reports (30, 31), we did not find p28-Omp19 expression in either wild-type or in the asRNA mutant when cultured in ISE6 tick cells (Fig 3B).

**Figure 3.**
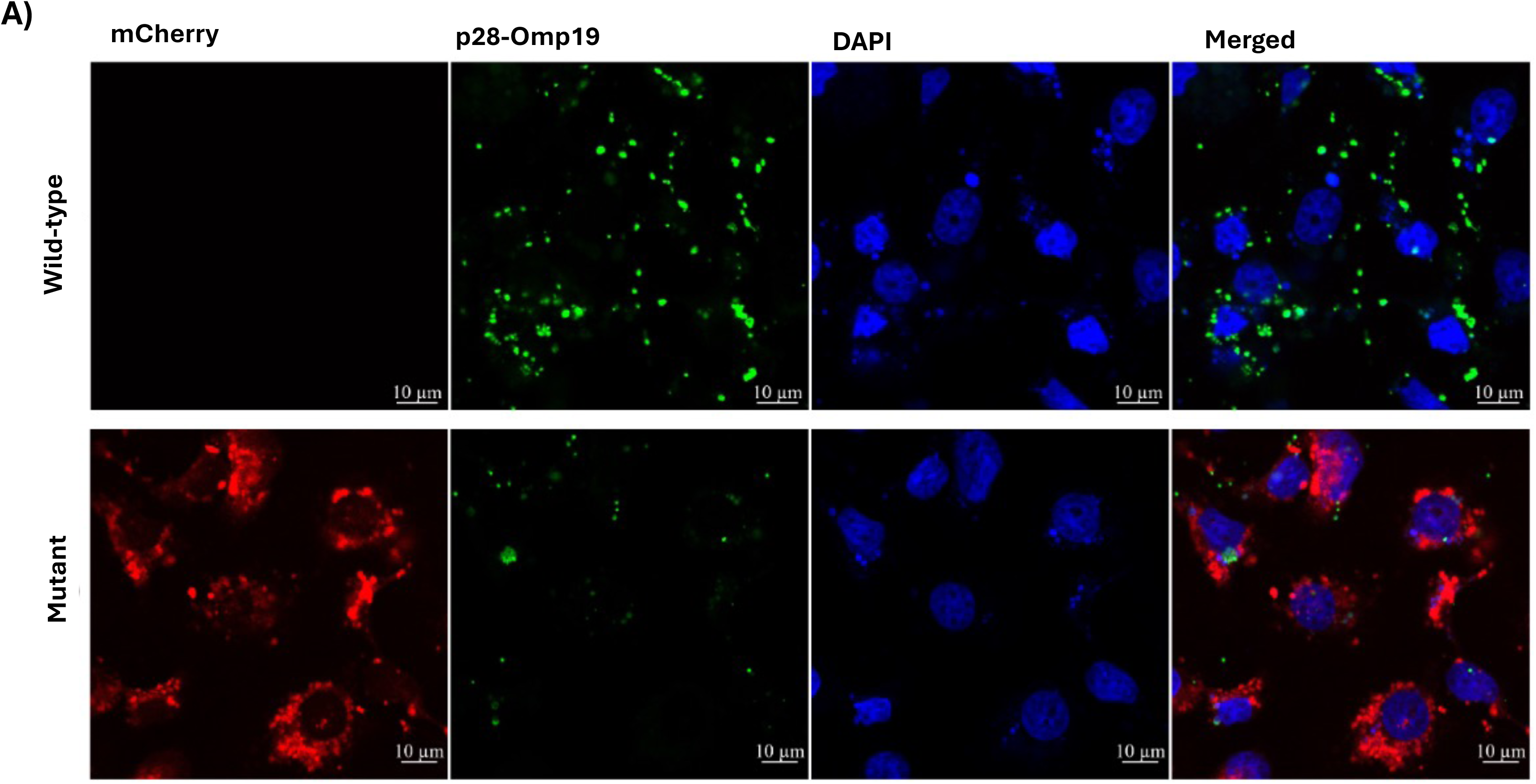
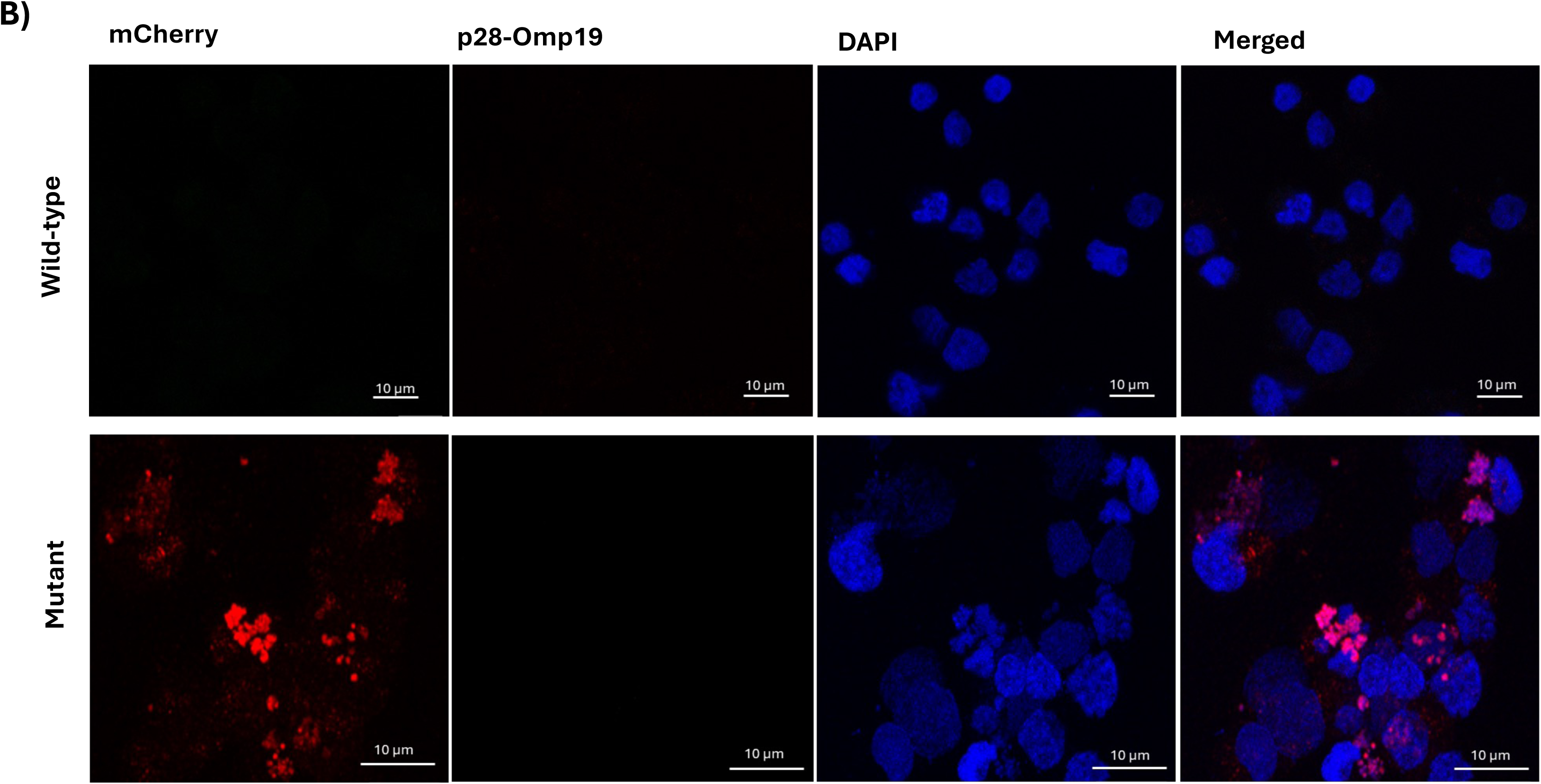
The impact of the *E. chaffeensis ECH_1143* antisense mutation on p28-Omp19 protein expression evaluated in different host cells by confocal microscopy. A) The mutation resulted in a significant reduction of p28-Omp19 expression in the macrophage cell line compared with the wild-type infection. B) P28-Omp19 was not evident in either wild-type or mutant when cultured in the tick cell line. Canine macrophage cell line (DH82) and tick cell line (ISE6) were infected with host cell-free wild-type or mutant *E. chaffeensis* and monitored when infection reached to about 80% cells are infected. The infected cells were then immunostained using the monoclonal antibody specific to p28-Omp19 (mAb 18.1) (green), while the host and bacterial nuclei were stained with DAPI (blue). Mutant bacteria expressing the mCherry (red) fluorescent protein under the constitutively active *tuf* promoter aided in monitoring total bacterial infection.

#### Western blot analysis confirmed the reduction in p28-Omp19 expression in macrophage cultures

Total resolved protein profiles of asRNA mutant and wild-type *E. chaffeensis* organisms were very similar independent of growing in macrophage and tick cell cultures (Fig 4A and D). Western blot analysis using *E. chaffeensis*-infected dog polyclonal sera (pAb) also revealed no major notable differences for the mutant compared to wild-type (Fig 4B and E). Whereas similar Western blot analysis using p28-Omp19-specific mAb 18.1 (Fig 4C and F) revealed the presence of considerably less protein being expressed from the mutant-derived protein fraction when cultured in the macrophage cell line. The p28-Omp19 protein expression was not detected in the immunoblots for tick cell culture-derived *E. chaffeensis* total proteins independent of the mutant or wild-type (Fig 4F). We then assessed the changes in the protein expression profile over a period of four days in macrophage culture for the mutant and wild-type *E. chaffeensis*. Total protein profiles looked similar for the time points assessed for the mutant and wild-type (Fig 5A). The immunoblot analysis performed with the mAb 18.1 revealed a steady increase in the p28-Omp19 expression over time for the mutant and wild-type *E. chaffeensis* with substantially less of the protein detected in the mutant-derived resolved protein fraction. The differences were more pronounced for the protein fractions recovered from 72 h and 96 h time points (Fig 5B).

**Figure 4.**
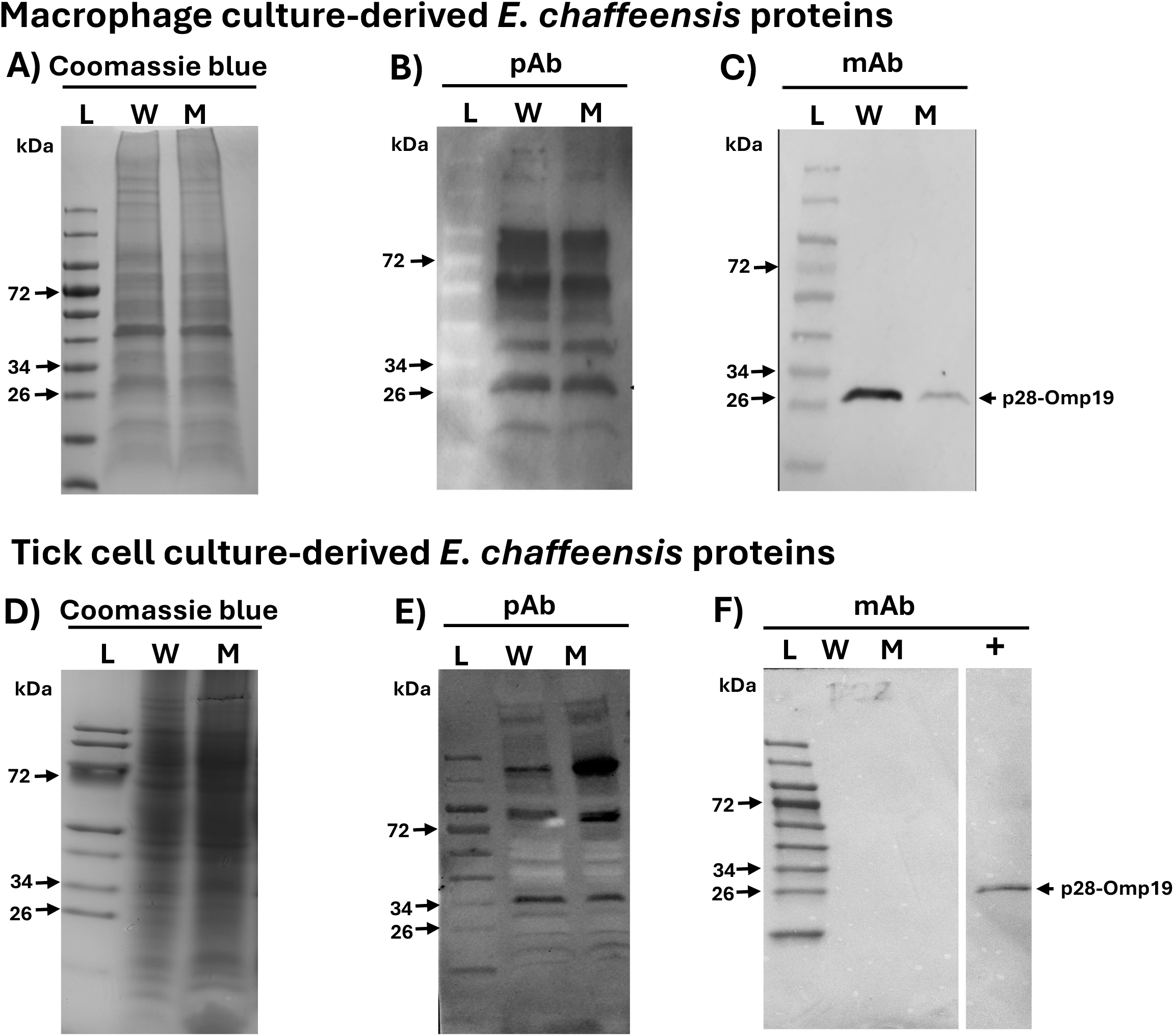
*E. chaffeensis ECH_1143* antisense mutation resulted in a reduction of p28-Omp19 protein expression in macrophage cultures (assessed by Western blotting). Macrophage and tick cell culture-derived *E. chaffeensis* wild-type and mutant lysates were subjected to PAGE and stained with Coomassie blue stain to visualize total proteins (3A and D). Western blotting of the bacterial lysates assessed using *E. chaffeensis*-infected dog polyclonal sera (3B and E) or with p28-Omp19-specific mAb (3C and F). L, protein ladder; W, wild-type; M, mutant; and +, positive control having *E. chaffeensis* protein extract from macrophage cultures.

**Figure 5.**
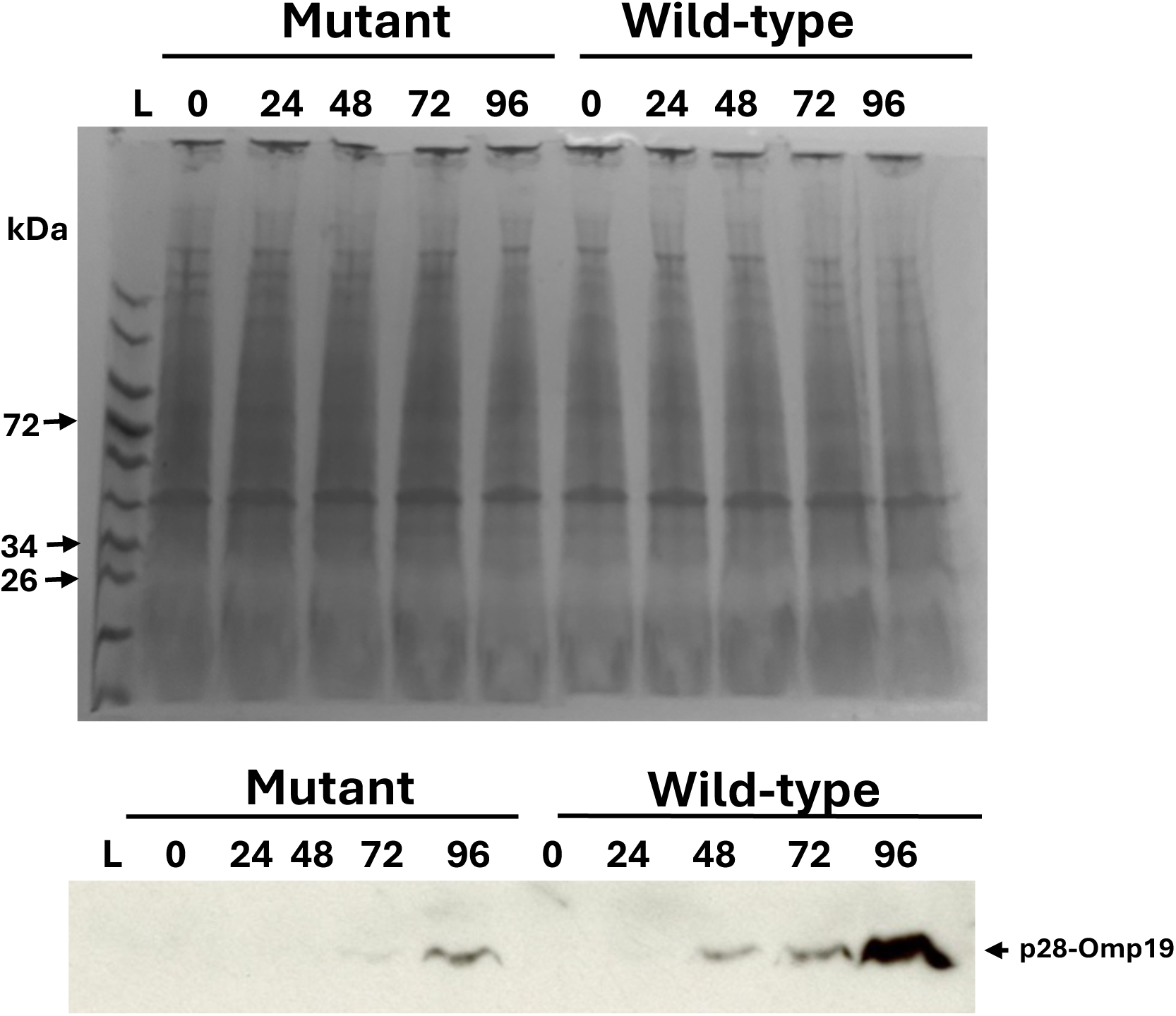
P28-Omp19 expression assessed over several days of growth of *ECH_1143* antisense mutant and wild-type *E. chaffeensis* cultured in the macrophage cell line. A) Wild-type and mutant lysates were subjected to PAGE and stained with Coomassie blue. B) Western blot analysis of lysates collected at different time points showed a relatively higher p28-Omp19 expression over time in the wild-type compared with that in the antisense mutant. L, protein ladder; the numbers above the blots refer to hours of culturing *E. chaffeensis* in macrophage cell line.

#### The asRNA mutant had attenuated growth in culture

We then assessed the impact of the mutation on *E. chaffeensis* growth in the macrophage and tick cell cultures. Infection in cultures was initiated with host cell-free asRNA mutant bacteria and its growth was monitored relative to another *E. chaffeensis* mutant similarly expressing mCherry but not having any gene defect in the genome to serve as a syngeneic control (Fig 6). The antisense mutant displayed attenuated growth rates compared to the growth of the control in both macrophage and tick cells; the altered growth for the asRNA mutant was more pronounced when cultured in the tick cell line compared to its growth in the macrophage cell line. Significant decline in the bacterial growth was evident for the mutant throughout the assessment period in tick cell culture beginning 12 h post initiation of the growth kinetics (Fig 6A). While the growth decline trend was evident for the mutant when cultured in the macrophage cell line, we observed high variation for the replicate samples despite triplicate samples assessed each time; a significant drop for the mutant’s growth in macrophage cell line was only evident for the last time point of assessment (Fig 6B).

**Figure 6.**
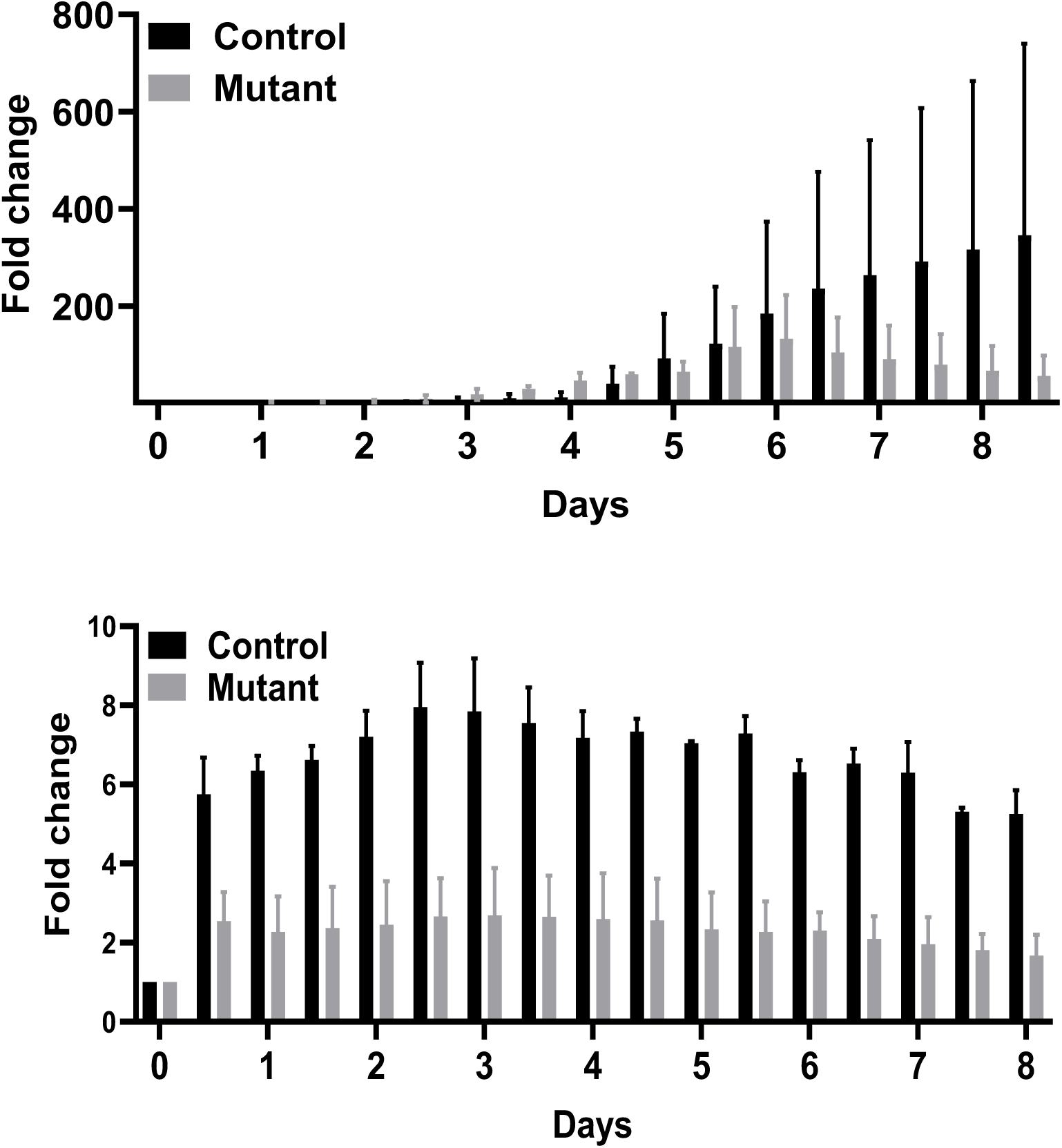
The continuous growth kinetics of the antisense mutant and another *E. chaffeensis* mutant not having any changes in the genome but expressing a targeted insertion expressing mCherry (control) were assessed. Infection in the host cells (macrophages and tick cells) was initiated with cell-free bacteria and growth was monitored over time using the number of mCherry fluorescent objects analyzed through confocal microscopy. Statistical significance was determined using two-way ANOVA with multiple comparisons. While macrophage-derived *E. chaffeensis* showed a clear trend of decline in the bacterial growth, statistically significant difference was only observed for the mutant for the last time point of assessment (*p <0.05) due to high variability for the replicate sample analysis. The tick cell-derived *E. chaffeensis* growth differed significantly from the mutant at all time points of assessment beginning from 12 h (***p <0.0002) independent of the replicate samples assessed.

### The asRNA mutation resulted in changes to the RNA expression from a paralog, p28-Omp14

Following confirming the knockdown of p28-Omp19 expression, we assessed if the reduction in the protein expression correlates with decrease of transcript level from *ECH_1143*. We further examined how the asRNA mutation impacted transcription from a paralog gene *ECH_1136* encoding for p28-Omp14 protein. Protein and RNA expression from the two paralogs are known to differ *in vitro* and *in vivo* with p28-Omp19 expression being higher when the pathogen replicates in macrophages but not in tick cells, while the protein and mRNA synthesized from *ECH_1136* (p28-Omp14) are predominantly detected in *E. chaffeensis* grown in tick cells (30, 31, 33). RNA expression, assessed by qRT-PCR, revealed very similar transcript levels for the *ECH_1143* transcript for the mutant and wild-type in the macrophage culture although time-dependent changes in the levels of transcription were evident (Fig 7A, left panel). Similarly, transcript levels did not significantly differ for *ECH_1136* in macrophage cultures except for the 48-h time point where expression was significantly decreased in the mutant compared to wild-type *E. chaffeensis* (Fig 7A, right panel). Similar assessment in tick cell cultures revealed notable reduction in the mRNA levels of *ECH_1143* for the mutant with significant decrease in transcript levels observed for the later time points (72 h and 96 h) (Fig 7B, left panel). The mRNA levels for *ECH_1136* for the mutant were significantly higher for all time points of assessment compared to wild-type (Fig 7B, right panel).

**Figure 7.**
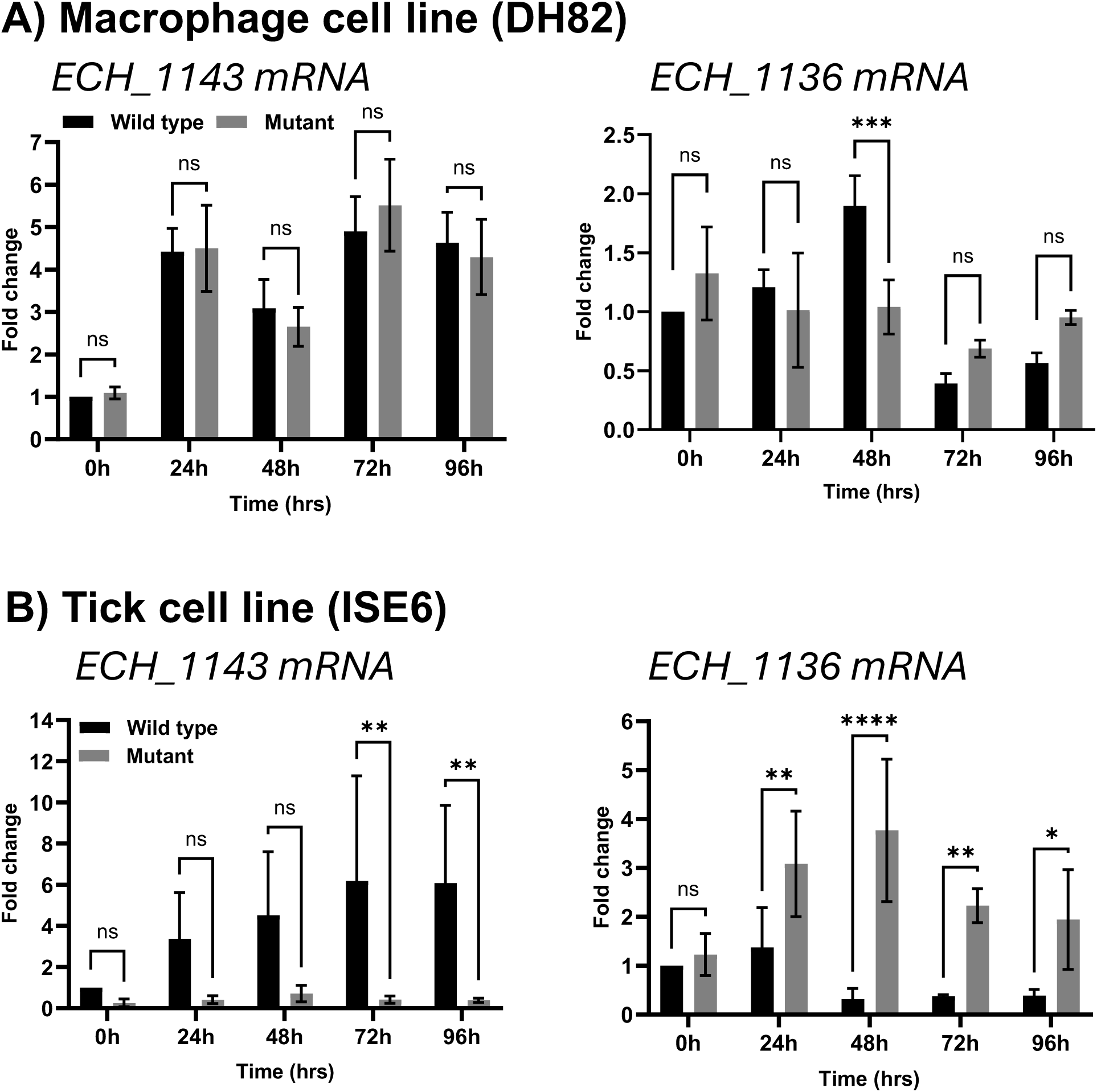
Transcription changes from genes *ECH_1143* and *ECH_1136* encoding for the proteins p28-Omp19 and p28-Omp14, respectively were assessed by qRT-PCR for the mutant and wild-type *E. chaffeensis* cultured in macrophage and tick cell cultures. Total RNA from synchronously cultured mutant and wild-type *E. chaffeensis* at different time points following infection of host cells were isolated. Real-time qRT-PCR analysis was performed to assess the changes in the mRNA levels from the two genes. Relative gene expression values are presented as fold changes compared to the wild type at 0 h post-infection. Data are shown as mean values ± standard deviations from three independent experiments and each experiment was also performed in triplicate. Statistical significance was determined using a two-way analysis of variance (ANOVA) followed by Tukey’s multiple-comparison test (*p < 0.05; **p < 0.01).

### Ribonuclease III (RNAse III) expression in *E. chaffeensis* is significantly higher during its growth in tick cells

We predicted that the observed reduction in p28-Omp19 expression as a result of the asRNA mutation correlates to reduction of *ECH_1143* mRNA when cultured in macrophages but we did not observe this. One of the mechanisms of reduction in protein expression in bacteria with sense and antisense RNA duplex formation is to sterically block translation initiation without degrading the duplex (36–38). This can occur when RNAse III expression is less (39). Bacterial RNase III regulates gene expression and mRNA turnover by degrading double-stranded RNA molecules (dsRNA) (34, 35). We, therefore, evaluated if RNAse III gene (*ECH_1054*) expression levels differ in wild-type *E. chaffeensis* when cultured in macrophage and tick cell lines (Fig 8). We detected higher RNAse III transcript levels for the bacteria in tick cell cultures throughout the four-day assessment period compared to macrophage culture with significant differences were evident at 24 and 72 h time points. The differences in RNase III expression may contribute to altered processing of asRNA-mRNA hybrids when the mutant *E. chaffeensis* replicates in tick cells vs. macrophage cultures.

**Figure 8.**
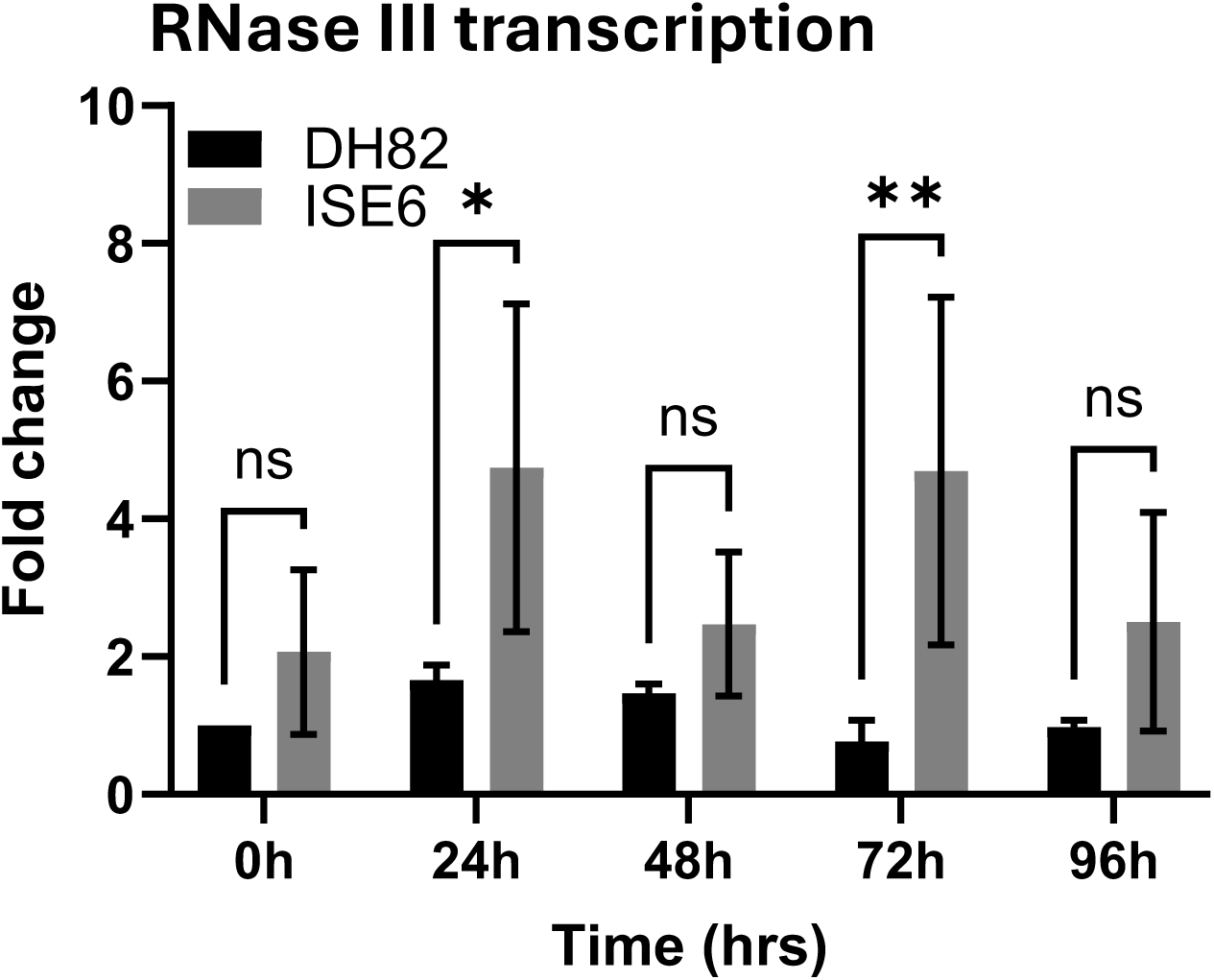
The RNase III transcript levels encoded from *ECH_1054* for wild-type *E. chaffeensis* grown in the macrophage and tick cells were measured. Total RNA isolated from synchronously cultured wild-type *E. chaffeensis* following different time points in culture. Real-time qRT-PCR analysis was performed to assess the expression levels of RNase III transcripts. Expression values are presented as fold changes relative to the 0 h time point. Data are shown as mean values ± standard deviations from three independent experiments, each performed in triplicate. Statistical significance was determined using a two-way analysis of variance (ANOVA) followed by Tukey’s multiple-comparison test (*p < 0.05; **p < 0.01).

## DISCUSSION

In the current study, we developed a novel targeted mutagenesis method in *E. chaffeensis* enabling the generation of *trans*-acting asRNA. The novelty of the method stems from expressing asRNA driven from the duplicated gene promoter involved in transcribing the sense mRNA thus providing a simultaneous expression of asRNA with sense mRNA leading to an effective means to downregulate the targeted gene expression. Further, when designing the mutational construct, we ensured that the mutation we retained the integrity of the genomic sequences unaltered at a distal genomic site where the mutation was introduced. We demonstrated the proof of concept in *E. chaffeensis* involved in generating asRNA expression by means of allelic exchange mutagenesis. This is the first description of an antisense knockdown system for a rickettsial pathogen, and we believe that the method will be broadly applicable for other bacterial pathogens.

Non-coding RNAs regulate gene expression in a wide range of species (40–43). In bacteria, such RNAs encoded for asRNAs modulate protein expression post-transcriptionally by base-pairing interactions with target sense mRNAs (43, 44). Non-coding antisense oligonucleotide (ASO)-mediated knockdown of a gene expression is a research tool aiding investigations in various organisms (21, 45). However, such studies are yet to be described in any known rickettsial pathogens. Non-coding RNAs, including *cis*-acting and *trans-*acting RNAs naturally exist in *Rickettsia* species and may regulate gene expression (46–49), although much remains to be understood about their functional significance. The existence of noncoding asRNAs is also reported in *Anaplasma phagocytophilum* and *Wolbachia pipientis* (50, 51), while their functional roles also remain unclear. An ASO–based gene experimental knockdown relies on the use of a delivery system tailored to an organism under investigation (21, 22, 52). In a plasmid-based delivery system, antisense DNA sequences are cloned downstream of an unrelated promoter to allow transcription of asRNA following such plasmid’s introduction into bacteria which results in the translation knockdown with or without promoting the sense RNA degradation (24, 53, 54). Traditional bacteria mutagenesis methods are challenging to be used for obligate, intracellular bacteria. Their dependence on the host cell for survival/replication makes working with the bacteria outside the host environment rather challenging and delicate. Additionally, the absence of natural plasmids harbored within the genomes of Anaplasmataceae family pathogens, including for *E. chaffeensis*, makes plasmid-based studies refractory (8). Transfection of chemically synthesized short antisense oligonucleotides may be used as an alternate tool for knocking down a gene expression (21, 55). This is also not a practical option as such oligonucleotides are highly prone to nuclease degradation. Recent studies described the application of transfected stable synthetic polymers, with characteristics similar to RNA and DNA, known as peptide nucleic acids (PNAs) to inhibit gene expression in *E. chaffeensis* (56). While this is a valuable alternative, the method is limited to transiently reducing protein expression (57).

The genetic modification method we reported in the current study involved creating a mutation at a distal genomic site to express asRNA complementary to the partial coding mRNA from *ECH_1143*. In this approach, we engineered the mutation downstream to the *ECH_0230* coding sequence, a region previously demonstrated as amendable for mutation (8). Further, we carefully designed the mutational construct to ensure that the integrity of the genomic sequences at the mutation site are not altered; the mutational deign included duplicating a segment of DNA 3’ to *ECH_0230* coding region to ensure that we do not delete any noncoding and possible regulatory sequences involved in the transcription from it or the gene positioned 3’ to it (*ECH_0232*). The mutational construct design included using the entire gene promoter segment located upstream to *ECH_1143* and then inserting 209-base pair open reading frame sequence from its initiation codon engineered in the reverse and complementary orientation. Our engineered mutagenesis cassette included the promoter segment of the target gene (*ECH_1143*) for facilitating the asRNA expression concurrently with the sense RNA synthesis. Synchronous expression of sense and antisense mRNAs is expected to result in an immediate and effective reduction in the protein output from the target gene. The data presented in the current study confirms the protein expression knockdown evidenced by two independent methods: confocal microscopy and Western blot analysis.

Protein expression from *ECH_1143* is host cell-specific both *in vitro* and *in vivo* (30, 31, 33). Prior studies demonstrated that it is a highly expressed immunogenic protein in *E. chaffeensis* during its growth in macrophage cultures, while it is not known to be expressed during its replication in tick cell cultures (33). *ECH_1136* is one of the 22 gene paralogs, including *ECH_1143*, existing in *E. chaffeensis* genome and it’s encoded p28-Omp14 protein is the only known expressed protein during the bacterial replication in tick cell environment (33). Similarly, the host-specific differential expression from *ECH_1143* and *ECH_1136* is documented *in vivo* (30); both paralogs are known to be expressed during infection of a vertebrate host, whereas *ECH_1136* is only expressed during infection in ticks (30, 31, 33). The data presented in the current study demonstrate that the novel antisense mutation caused a substantial reduction in protein expression from *ECH_1143* when the pathogen was cultured in a macrophage cell line. Furthermore, we also demonstrated that protein expression was undetectable for it during replication in tick cells independent of wild-type or mutated *E. chaffeensis* despite the presence of mRNA. These data are in agreement with the prior reports of the absence of p28-Omp19 expression in tick cell cultures (33).

We observed the presence of *ECH_1143* transcript in tick cells for wild-type despite no detectible protein expression and the asRNA mutation reduced the transcript levels with increased transcription from a paralog, *ECH_1136*. Whereas the transcript levels remained similar for wild-type and the mutant in the macrophage culture for *ECH_1143* and similarly for *ECH_1136*, except for the 48 h time point assessment. Interestingly the mutation negatively impacted *E. chaffeensis* growth in both macrophage and tick cell backgrounds. The data suggest that the antisense mutation knockdown mRNA in tick cells may have a complementary effect in enhancing the transcription of *ECH_1136.* It is unclear why p28-Omp19 expression was not evident for the pathogen during its growth in tick cells. One possible explanation is that the protein expression is too low to be detected by the methods we used: confocal and Western blot methods. While warranting additional investigations, the data also suggest that the bacterial gene regulation is heavily impacted by its replication in vertebrate and invertebrate host cell backgrounds. Indeed, prior published research reported host-specific differences in *E. chaffeensis* gene expression (33). The lack of RNA expression changes in the mutant compared with wild-type while causing the protein expression knockdown suggests that asRNA only blocked translation but not mRNA degradation in macrophage cultures. Conversely, *ECH_1143* mRNA expression reduction was evident for the mutant in tick cells, and the data suggest that asRNA may have induced sense mRNA breakdown during growth in tick cell cultures. Binding of asRNA to target mRNA blocks translation by inhibiting ribosome access; alternatively, it can promote RNase III-mediated degradation of the duplex, and both mechanisms lead to the protein synthesis reduction (58). Therefore, the interactions between sense RNA and asRNA do not always lead to mRNA degradation (38). Instead, the RNA duplex may create a specific processing site, which can result in a translationally inactive mRNA or produce a mature and stabilized form of duplex mRNA (38). The asRNA of *ECH_1143* when bound within a window of 70 codons from the start codon may have likely inhibited ribosome binding and decreased p28-Omp19 expression without inducing degradation of the sense RNA during the mutant’s replication in the macrophage cell line.

In *E. coli*, RNase III mediates the breakdown of sense-antisense hybrids (59, 60). Our results demonstrate that the *E. chaffeensis* RNase III transcription differed for the pathogen when grown in macrophage and tick cell cultures. RNase III is significantly higher during *E. chaffeensis* replication in tick cell cultures compared to its replication in macrophage cultures. These data were similar to the previously reported genome-wide *E. chaffeensis* gene expression assessment which also suggests that mRNA levels of RNase III are lower for the organisms cultured in human monocytic cells (THP-1) compared to tick cells (ISE6) (61). Thus, it is likely that having higher RNAse III expression in tick cell cultured organisms may have promoted the breakdown of sense and antisense mRNA duplexes. The data also suggest that *E. chaffeensis* employs distinct post-transcriptional regulatory mechanisms in macrophage and tick cell environments.

In summary, the novel mutational approach is established for the first time to generate a protein expression knockdown for an obligate intracellular bacterial pathogen, *E. chaffeensis.* It is a pertinent alternative for investigating genes which are likely essential for pathogens having reduced genomes. The method is unique in allowing simultaneous asRNA expression as both the sense and antisense RNAs are transcribed from the same gene promoter, as the mutation included the duplicated gene promoter that belongs to the same gene. This innovative molecular tool will greatly facilitate research in investigating essential pathogen genes and is likely applicable to other bacterial pathogens having reduced genomes, such as those within the order of Rickettsiales and possibly for other important pathogenic microorganisms belong to the genera *Borrelia, Chlamydia and Coxiella*.

## MATERIALS AND METHODS

### In vitro cultivation of E. chaffeensis

*E. chaffeensis* Arkansas isolate was continuously cultivated in the *Ixodes scapularis* embryonic cell line (ISE6) and in the canine macrophage cell line (DH82) (macrophage) as previously described (46).

### Construction for homologous recombination plasmids and segments

The primers designed for use in generating the recombinant plasmid are listed in supplementary Table S1. All PCR experiments for preparing the recombinant plasmid construct were performed using Q5 High-Fidelity DNA Polymerase (New England Biolabs, Ipswich, Massachusetts). The 5′ *E. chaffeensis* left homology arm (L) with genome coordinates of the amplified segment spanning from 218,537 to 219,618 and 3′ right homology arm (R) with genome coordinates of the segment are 219,083 to 220,133 were amplified (GenBank accession no. CP000236). The L segment spans from part of *ECH_0230* open reading frame (ORF) and the entire noncoding region located 3’ to the ORF and the R segment also contained the same entire noncoding region located 3’ to the *ECH_0230* ORF and the partial ORF belong to *ECH_0232* (3’ portion of the ORF). The complete *ECH_1143* promoter segment (genome coordinates:1,163,653 to 1,163,967) and the first 209 base pair (bp) ORF sequence of *ECH1143* (genome coordinates: 1,164,176 to 1,163,968) were independently amplified using wild-type *E. chaffeensis* genomic DNA as the template. The *ECH_1143* promoter segment and the 209 bp *ECH_1143* ORF cloned in reverse orientation into a plasmid; the assembled fragment was referred as AS. The *tuf* promoter (T) along with the mCherry (M) and gentamicin resistance protein coding sequences (G) were obtained from the AM581-KO-tuf-mCherry-Gent plasmid which we reported previously (10). All these fragments were cloned in into the pGGA plasmid (New England Biolabs, Ipswich, MA) in the following order: L, T, M, G, AS, and R, using a Golden Gate Assembly kit (New England Biolabs, Ipswich, MA). Recombinant plasmid was transformed into *E. coli* DH5α, and plasmid DNA was recovered. Correct assembly of the recombinant plasmid DNA segments was confirmed by Sanger sequencing with T7 and SP6 primers (Integrated DNA Technologies, Coralville, IA) which anneal upstream and downstream of the recombinant DNA segment in the pGGA plasmid. The final construct was named as pHR-*ECH_1143*-Antisense. This plasmid served as the template for amplification of the allelic exchange fragment containing the recombinant segment spanning from L to R in the plasmid. PCR products were then purified with the QIAquick PCR Purification Kit (Qiagen, Germantown, MD) prior to using for the mutation generation.

### Electroporation of *E. chaffeensis* with the linear DNA fragments and the selection of the mutant

Host cell-free *E. chaffeensis* organisms were recovered as previously described in (11). The linear purified DNA fragments (3 µg) generated above from the pHR-*ECH_1143-*Antisense plasmid were added to the host cell-free *E. chaffeensis* organisms in 45 µl volume, mixed gently and transferred to a 1 mm electroporation cuvette (Bio-Rad Laboratories, Hercules, CA), placed in ice for 15 min prior to subjecting to electroporation at 2,000 volts, 25 µF and 400 Ω setting (Gene Pulser Xcell™, Bio-Rad Laboratories, Hercules, CA). The electroporated cells were transferred to a micro centrifuge tube containing 0.5 ml of stock FBS and 1 ml of uninfected ∼1x10^6^ ISE6 cell suspension in the tick cell culture infection media, then centrifuged at 5,000 g for 5 min, and incubated at room temperature for 15 min. The cell suspension was transferred to 5 ml culture media, transferred to a T25 flask having a confluent ISE6 cell monolayer, and incubated for 24 h at 34 °C in a humidified incubator. Subsequently, 80 µg/ml of gentamicin was added to the culture medium after 24 h, and cultures were monitored for several weeks until mCherry expressing *E. chaffeensis* infection was detected and the wild type was not detected.

### Confirming the presence of *E. chaffeensis* mutants

The *E. chaffeensis* culture resistant to the presence of gentamicin in media was screened to confirm the mutation. Genomic DNA recovered from the mutant cultures was used to perform two different PCR assays. PCR I was targeted to define the clonal purity of the mutant where the primers used for the assay were targeted to the genomic regions upstream and downstream to the allelic exchange insertion site. PCR II was targeted to the genomic region 5’ to the allelic exchange site and to the insertion specific DNA; PCR primers for both PCR I and II were listed in the Supplementary Table S1. The PCR products were resolved on a 0.9% agarose gel to check for the presence of specific predicted amplicons. Additionally, whole genome sequencing was performed using mutant genomic DNA on Nanopore sequencing platform (Plasmidsaurus Inc., Kentucky, USA). To remove Nanopore adapters and to perform quality filtering, Porechop v.024 (62) and Filtlong v.0.2.1 (https://github.com/rrwick/Filtlong) was used with default settings. The remaining high-quality reads were mapped to the *E. chaffeensis* isolate reference genome (CP000236) and the insert construct (mCherry gene, gentamicin gene, the promoter of *ECH_1143* and antisense sequence of *ECH_1143*) using Minimap2 v.2.28 (63). Mapped reads were extracted via samtools v.1.21 (64) and assembled *de novo* using Flye v.2.9.6 (65). The Artemis genome browser and annotation tool is used to visualize the sequence data (66).

### P28-Omp protein expression in the mutant and wild-type *E. chaffeensis* assessed by confocal microscopy analysis

At approximately 80% infection*, E. chaffeensis* wild-type and mutant-infected macrophage and tick cell cultures were seeded in a 24-well plate (containing a glass slip) or four-chambered slides at a density of 1 × 10^5^ cells per well, respectively. The supernatant medium was removed, and the cells were washed three times with phosphate buffered saline (PBS). Cells were fixed with 4% paraformaldehyde solution for 10 min at room temperature. Cells were then permeabilized with 0.2% Triton X-100 in PBS (PBST) for 15 min at room temperature, then washed three times with PBS. Cells were blocked with 5% bovine serum albumin (BSA) buffer for 2 h at room temperature, then incubated with the primary antibody, p28-Omp19-specific monoclonal antibody, mAb 18.1, raised in mice (1:200 or 400) (32), at 4°C overnight. Cells were washed three times with PBS or 1X PBST. Cells were then incubated with Alexa Fluor 488-conjugated Goat Anti-mouse IgG (H+L) antibody (1:2000) for 2 h at room temperature or 45 min-1 h at 37 4°C with cells protected from light. After three washes with PBST, cells were stained with DAPI solution for 10 min at room temperature or ProLong Diamond antifade mountant with DAPI (Thermo Fisher Scientific, Waltham, MA) for 5 min at room temperature. Images were captured using an LSM 880NLO laser (Carl Zeiss, Germany) and MP confocal microscopes and further processed in ImageJ.

### Protein expression over time in *E. chaffeensis* wild-type and mutant when cultured in the macrophage or tick cell lines and assessed by Western blot analysis

Macrophage and tick cell culture-derived *E. chaffeensis* wild-type and mutant were purified as previously described (33). Bacterial lysates were prepared, and total protein concentrations were measured using the bicinchoninic acid (BCA) protein assay as per the manufacturer’s recommendations (Thermo Fisher Scientific, Waltham, MA). Lysates were then subjected to Sodium Dodecyl Sulfate Polyacrylamide Gel Electrophoresis (SDS-PAGE) and either stained with Coomassie blue stain to visualize the separated proteins or transferred to a polyvinylidene difluoride (PVDF) membrane for use in Western blot analysis. The membranes were blocked with 5% non-fat dry milk in Tris-buffered saline with 0.1% Tween 20 (TBST) for 1 h at room temperature, then incubated with mAb 18.1 (1:1000) specific to detecting *E. chaffeensis* p28-Omp19 or polyclonal sera from a *E. chaffeensis* wild-type-infected dog (1:500) (70, 71) at 4°C overnight. After three washes with TBST for 5 min each, membranes were incubated with horseradish peroxidase (HRP)–conjugated Goat anti-mouse or anti-dog at a 1:4000 dilution in TBST for 1 h at room temperature. Protein bands were detected using an enhanced chemiluminescence (ECL) Western blot detection kit (Thermo Fisher Scientific, Waltham, MA) according to the manufacturer’s instructions. The images of Western blots were capture using iBright Imaging Systems (Thermo Fisher Scientific, Waltham, MA).

### Antisense mutant growth assessed by real-time live cell confocal image analysis

To measure the growth rate of the antisense mutant real-time, we used a BioTek Cytation C10 Confocal Imaging Reader (Agilent, USA). To facilitate the growth comparison requires to include a wild-type equivalent *E. chaffeensis* that is similarly expressing mCherry protein. Therefore, we used another mutant *E. chaffeensis* that is having no genetic defects other than that it expresses mCherry protein constitutively expressed from *tuf* promoter as in the antisense mutant. (This mutant was generated as part of another study; our unpublished results). We grew the two versions of *E. chaffeensis* to measure fluorescent units over a period of several days of continuous culture; the assay was performed in triplicate wells, and it was repeated three independent times. The measurements were taken once every four hours. Average values for the fluorescent measurements were plotted with a half-day intervals to define the the growth of the organisms in macrophage and tick cells.

### RNA analysis by TaqMan probe-based real-time quantitative RT-PCR (qRT-PCR) to define the impact of the antisense mutation

For synchronous growth, macrophage and tick cell cultures at a concentration of 1x10^5^ cells per milliliter were plated in each well of a six-well plate. When the cell confluency reached approximately 70-80%, each well was inoculated with host-cell-free mutant or wild-type *E. chaffeensis*. To allow bacterial internalization, the cultures were incubated for 2-3 h following which the media was removed and replaced with fresh media. This served as zero time point and cells from one well each were harvested at zero time point and similarly after every 24 h from then on for four days. Total RNA from each harvested sample was purified using Tri-reagent total RNA isolation kit method (Sigma-Aldrich, St. Louis, MO). DNA contamination from the purified RNAs was eliminated by treating with RQ1 DNase (Promega, Madison, WI). RNA samples were equalized relative to 16S rRNA expression as we described earlier in (72). Primers and probes for the TaqMan probe-based RT-PCR assays for determining the RNA expression from *ECH_1136*, *ECH_1143*, and for the RNase III gene (*ECH_1054*) were listed in the Supplementary Table S1 which were used to evaluate changes in the mRNA levels for each gene for all harvested RNAs. The fold differences relative to zero time point were calculated by 2^(-Delta Delta Ct) method (73). The analysis was performed three independent times and represented the mean values of the three experiments.

### Statistical analysis

All data were analyzed by a two-way analysis of variance (ANOVA) with SPSS 25.0 (IBM SPSS Inc., Chicago, IL) and using GraphPad Prism 12 software (GraphPad Software, La Jolla, CA, USA). All data are presented as the means ± standard deviation (SD). *P* < 0.05 and *P* < 0.01 represented the significant and highly significant difference, respectively.

## Acknowledgements

This research was supported by the PHS grants R01AI070908 and R01AI182519 from the National Institute of Allergy and Infectious Diseases, National Institutes of Health, USA.

